# Generalized displacement of DNA- and RNA-binding factors mediates the toxicity of arginine-rich cell-penetrating peptides

**DOI:** 10.1101/441808

**Authors:** V. Lafarga, O. Sirozh, I. Díaz-López, M. Hisaoka, E. Zarzuela, J. Boskovic, B. Jovanovic, R. Fernandez-Leiro, J. Muñoz, G. Stoecklin, I. Ventoso, O. Fernandez-Capetillo

## Abstract

Due to their capability to transport chemicals or proteins into target cells, cell-penetrating peptides (CPPs) are being developed as therapy delivery tools. However, and despite their interesting properties, arginine-rich CPPs often show toxicity for reasons that remain poorly understood. Using a (PR)n dipeptide repeat that has been linked to amyotrophic-lateral sclerosis (ALS) as a model of an arginine-rich CPP, we here show that the presence of (PR)n leads to a generalized displacement of RNA- and DNA-binding proteins from chromatin and mRNA. Accordingly, any reaction involving nucleic acids such as RNA transcription, translation, splicing and degradation or DNA replication and repair are impaired by the presence of the CPP. Interestingly, the effects of (PR)n are fully mimicked by PROTAMINE, a small arginine-rich protein that displaces histones from chromatin during spermatogenesis. We propose that widespread coating of nucleic acids and consequent displacement of RNA- and DNA-binding factors from chromatin and mRNA accounts for the toxicity of arginine-rich CPPs, including those that have been recently associated to the onset of ALS.

## INTRODUCTION

Efforts to develop methods to introduce nucleic acids or proteins into cells started in the mid XX^th^ century, with the realization that alkaline conditions favored the infectivity of poliovirus RNA in HeLa cells (Sprunt et al., 1959). Soon thereafter, it was found that the addition of basic proteins such as histones or PROTAMINE also enhanced the uptake of RNA or ALBUMIN by tumor cells in culture (Ryser and Hancock, 1965; Smull and Ludwig, 1962). These early studies already noted that arginine-rich histone fractions were more efficient than lysine-rich ones in stimulating the cellular uptake of proteins. In fact, several factors with arginine-rich domains such as fibroblast growth factor (bFGF), pancreatic Ribonuclease A (RNase A) or the cytokine midkine (MK) are known to freely enter into cells (reviewed in (Fuchs and Raines, 2006)). The field of protein transfection regained attention in 1988 when 2 groups independently discovered that the HIV-1 trans-activator protein TAT could enter cells, accumulate at nuclei and nucleoli and be functional in trans-activating HIV-1 RNA transcription (Frankel and Pabo, 1988; Green and Loewenstein, 1988). The arginine-rich dodecapeptide GRKKRRQRRRPQ was later defined as the minimum functional sequence from TAT, which when fused to any given protein enables its cellular uptake (Park et al., 2002). Today, cell penetrating peptides (CPPs) are intensively being developed due to their capacity to facilitate the uptake of nucleic acids, small molecules, proteins or even viruses (reviewed in (Borrelli et al., 2018)).

Despite the interesting properties of CPPs, there are also concerns on their toxicity, particularly for those that are arginine-rich. For instance, and despite its clinical use as an antidote of the anticoagulant heparin, PROTAMINE has important side-effects which are driving the search for alternative heparin antidotes (Sokolowska et al., 2016). In addition, HIV-1 TAT is neurotoxic *in vitro*, an observation that has been proposed as a potential explanation for AIDS-associated neurodegeneration (King et al., 2006). Interestingly, studies done in the last decade have revealed that the pathogenicity of the most frequent mutation found in patients of amyotrophic lateral sclerosis (ALS) and frontotemporal dementia (FTD), which occurs at a gene named *C9ORF72*, might also be related to the production of arginine-rich CPPs. *C9ORF72* mutations involve the expansion of a GGGGCC hexanucleotide within the first intron of the gene, which is amplified to hundreds or even thousands of copies in ALS/FTD patients (DeJesus-Hernandez et al., 2011; Renton et al., 2011). Through repeat-associated non-AUG (RAN) translation sense and antisense GGGGCC expansions are translated into several dipeptide-repeats (DPR), including (PR)n and (GR)n (Ash et al., 2013; Mori et al., 2013; Zu et al., 2013). Significantly, synthetic (PR)n and (GR)n peptides added to culture media enter cells, accumulate at nucleoli and kill cells, thus behaving as toxic CPPs (Kwon et al., 2014). Accordingly, the expression of these DPRs is toxic in cells and animal models (Chew et al., 2015; Kwon et al., 2014; Mizielinska et al., 2014; Stopford et al., 2017; Swaminathan et al., 2018; Wen et al., 2014; Zhang et al., 2018). While the toxicity of arginine-rich CPPs seems to underlie several pathologies, the mechanisms by which this occurs remain unknown. By using the (PR)n DPR found in patients of ALS/FTD as a model of an arginine-rich CPP, we here show that its presence leads to a generalized displacement of RNA- and DNA-binding factors from chromatin and RNA, explaining its widespread effects on reactions involving nucleic acids.

## RESULTS

### (PR)_20_ peptides impair the assembly of 80S ribosomal particles on mRNA

Proteomic and genetic studies have revealed that arginine-rich DPRs arising from C9ORF72 mutations have a particular impact on mRNA translation (Chai and Gitler, 2018; Kanekura et al., 2016; Lopez-Gonzalez et al., 2016; Zhang et al., 2018). To further investigate how these DPRs affect translation, we performed a proteomic characterization of ribosomes purified from HeLa cells exposed to synthetic (PR)_20_ peptides (**Fig. S1A**). Ribosome purification was done by immunoprecipitation of a stably expresed streptavidin-binding peptide (SBP)-tagged RPS9. The most prominent observation in ribosomes isolated from (PR)_20_-treated cells was a generalized decrease in the abundance of L ribosomal proteins (RPLs) from the 60S subunit (**Fig. S1B, Table S1**), which was not due to a reduction in the total levels of these factors (**Fig. S1C**). Since ribosome purification was performed by pulling down RPS9, a component of the 40S ribosomal subunit, the observed reduction in 60S factors suggested that the peptide could be impairing the assembly of 80S ribosomes from 40S and 60S subunits. Accordingly, ribosome profiling from HeLa cells treated with (PR)_20_ peptides revealed an accumulation of polysome halfmers (**Fig. S1D**), which is indicative of an incomplete assembly of 80S ribosomes at sites of translation initiation. Interestingly however, (PR)_20_ did not impair the *in vitro* assembly of 80S particles from purified 40S and 60S subunits in the absence of mRNA, as evaluated by electron microscopy, arguing that the peptide did not directly affect the components of the 40S or 60S subunits (**Fig. S1E,F**).

To further investigate how (PR)n impairs translation, we performed *in vitro* translation reactions in rabbit reticulocyte lysates. Consistent with previous work (Kanekura et al., 2016), *in vitro* translation of a luciferase mRNA was impaired by (PR)_20_ in a dose-dependent manner (**Fig. S1G,H**). The presence of polysome halfmers and the lower abundance of RPLs on ribosomes purified from (PR)_20_-treated cells suggested that the DPRs could be preventing the assembly of 40S and 60S subunits into 80S ribosomes by binding to mRNA at translation initiation sites. In fact, arginine-rich oligopeptides have very high biochemical affinity for RNA (Tan and Frankel, 1995) and ALS-associated arginine-rich DPRs specifically have been shown to bind to and form aggregates in the presence of RNA (Boeynaems et al., 2017; Kanekura et al., 2016; White et al., 2019). In support of this hypothesis, extending the length of the non-coding 5’UTR of the luciferase mRNA led to a further decrease in translation by (PR)_20_ (**Fig. S1I**), due to an increased probability of DPR-binding to the 5’UTR that could block initiation. Of note, the interaction with (PR)_20_ does not irreversibly alter RNA molecules, as mRNA purified from (PR)_20_-treated reticulocyte extracts was efficiently translated in a subsequent *in vitro* translation reaction performed in the absence of the DPR (**Fig. S1J**). Collectively, these experiments indicate that the mechanism by which (PR)_n_ peptides impair translation is due to interactions with mRNA that prevent the assembly of 40S and 60S subunits into 80S ribosomes on active polysomes.

### A general effect of (PR)_20_ peptides in reactions using DNA or RNA substrates

Besides translation, a number of cellular processes using RNA intermediates have been reported to be altered by *C9ORF72*-associated arginine-rich peptides including splicing (Kwon et al., 2014; Yin et al., 2017), rRNA biosynthesis (Kwon et al., 2014) and mRNA export (Rossi et al., 2015). While each of these effects have been proposed to be due to the binding of the DPRs to specific factors such as the splicing regulator U2 snRNP (Yin et al., 2017), we reasoned that if the effects of arginine-rich peptides on translation were due to RNA coating, any reaction using an RNA template should be similarly affected. Supporting this view, *in vitro* reactions using RNA substrates such as reverse transcription or RNase A-mediated RNA degradation were impaired by (PR)_20_ in a dose-dependent manner (**Fig. 1A,B**). Noteworthy, the fact that (PR)_20_ protects RNA from degradation can influence proteomic analyses aiming to identify proteins that bind to arginine-rich peptides, since this effect will prevent the exclusion of interactions that are indirectly mediated by RNA. In fact, proteomic studies looking for (PR)n interactors have consistently found an enrichment of RNA-binding factors (Kanekura et al., 2016; Lin et al., 2016; Yin et al., 2017). Regarding the cellular effects of (PR)_20_ related to RNA reactions, we could confirm an overall decrease in RNA biosynthesis (Kwon et al., 2014) (**Fig. 1C**) and an accumulation of mRNA in the nucleus indicative of deficient RNA export (Rossi et al., 2015) (**Fig. 1D**) in U2OS cells exposed to (PR)_20_. In addition, the presence of the peptide altered the nucleoplasmic distribution of Cajal Bodies, which are membrane-less organelles made up of RNA and proteins that are involved in various aspects of RNA metabolism including snRNA modifications, histone mRNA assembly or telomeric RNA processing (**Fig. 1E**) (Nizami et al., 2010). Noteworthy, the effects of (PR)_20_ are not restricted to cellular RNAs as the intracellular replication of Sindibis virus, which contains a single-stranded RNA genome, in Baby Hamster Kidney cells (BHK-21) was also impaired by the presence of (PR)_20_ (**Fig. 1F**), further supporting the notion that (PR)n DPRs have a general effect on cellular reactions using RNA substrates.

**Fig. 1.**
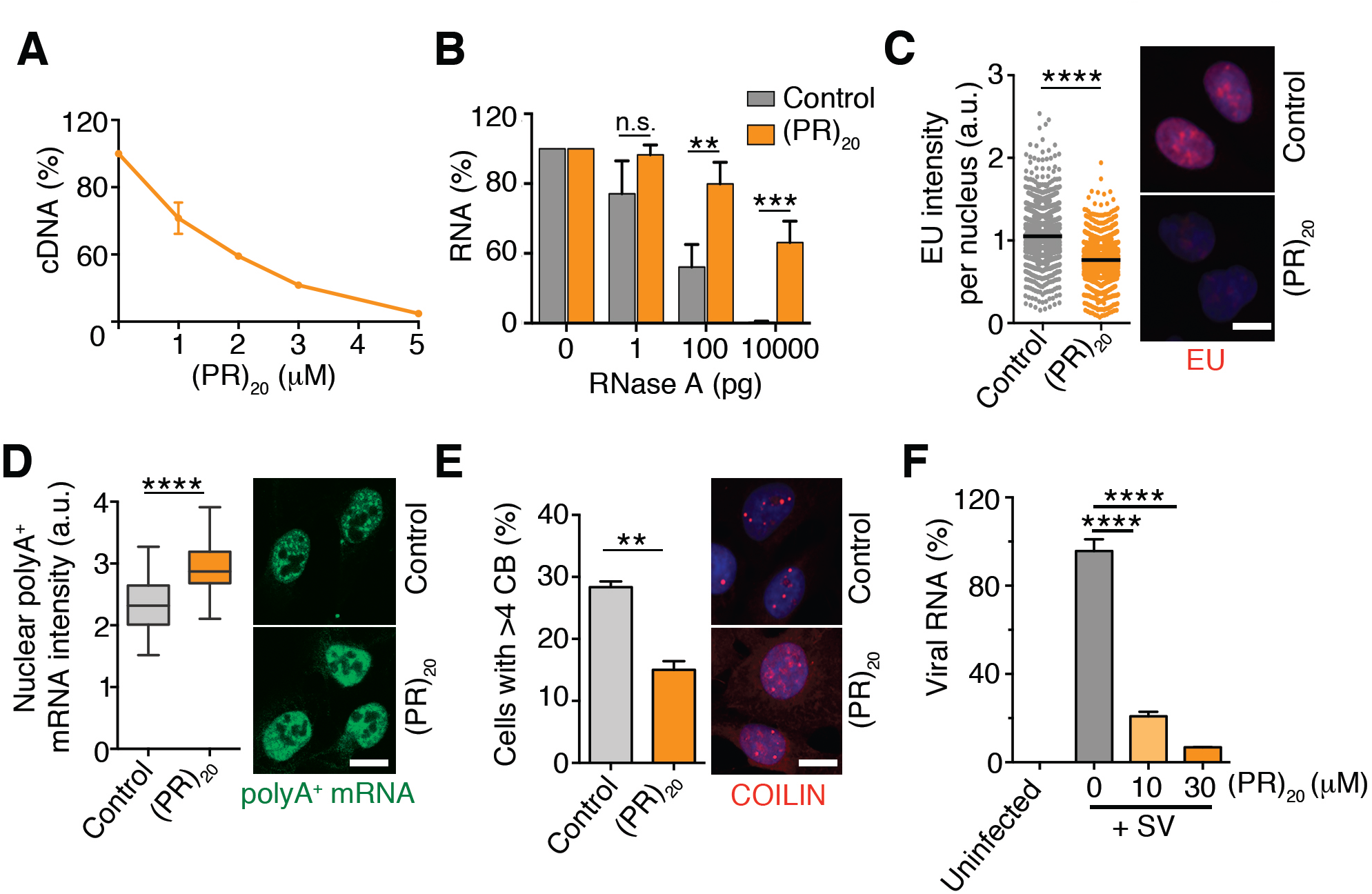
RNA-based reactions are collectively impaired by (PR)_20_ peptides. (**A**) Reverse transcription of RNA (500 ng) with oligo-dT in the presence of increasing doses of (PR)_20_. Data represent the fluorometric quantification of the resultant cDNA (n=2). (**B**) Percentage of RNA (1 µg) remaining after a 15’ digestion with increasing doses of RNase A in the presence or absence of (PR)_20_ (5 µM) (n=3). (**C, D**) High-throughput microscopy (HTM)-mediated analysis of 5-ethynyl-uridine (EU) (**C**) and polyA+ mRNA (**D**) levels per nucleus found in U2OS cells exposed to 10 µM (PR)_20_ for 16 h. EU was added 30’ prior to fixation. (**E**) Effect of (PR)_20_ (10 µM, 16 h) in the nucleoplasmic distribution of Cajal Bodies (CB) identified by immunofluorescence with anti-COILIN antibodies. Note that besides its impact on the number of CBs, (PR)_20_ also promotes a dispersed distribution of COILIN in the nucleoplasm. Representative images from these analyses (**C**-**E**) are provided in each case (right). Scale bar (white) indicates 2-5 µM. **F**, Accumulation of Sindibis virus (SV) genomic RNA in BHK-21 cells in the absence or presence of increasing doses of (PR)_20_. The amount of viral and cellular RNA was quantified from total RNA isolated 4 h after infection by quantitative real-time PCR (qRT-PCR) with specific primers. **, p<0.01; ***, p<0.001; ****, p<0.0001. See also **Fig. S1** for further analyses of the effect of (PR)_20_ in mRNA translation.

In addition to RNA, oligoarginine peptides are also known to bind avidly to DNA and promote its compaction (DeRouchey et al., 2013; Mascotti and Lohman, 1997; Tan and Frankel, 1995). Consistently, electrophoretic mobility assays (EMSA) revealed a similar affinity of (PR)_20_ for DNA and RNA in single- or double-stranded form (**Fig. 2A**). We thus explored the impact of (PR)_20_ in reactions using DNA substrates. In vitro, (PR)_20_ inhibited DNA polymerase chain reactions in a dose-dependent manner (**Fig. 2B,C**). When added to U2OS, the presence of (PR)_20_ reduced DNA replication rates (**Fig. 2D**) and impaired the repair of DNA breaks generated by ionizing radiation as measured with antibodies detecting the phosphorylated form of histone H2AX (γH2AX) (**Fig. 2E**). The efficiency of gene-deletion of an EGFP cDNA by CRISPR/Cas9 using a sgRNA against EGFP was also significantly affected by (PR)_20_ (**Fig. 2F**), although this could be mediated by effects of the peptide on DNA and/or RNA. In contrast to the widespread effect of (PR)_20_ in RNA- or DNA-based reactions, the presence of the DPR did not affect the efficiency of biochemical reactions not using nucleic acids such as an *in vitro* phosphatase assay (**Fig. S2A**) or the previously mentioned *in vitro* assembly of 80S ribosomal particles from purified 40S and 60S subunits (**Fig. S1E,F**). Of note, these effects seem to a large extent to be mediated by arginines, since even if a (PK)_20_ peptide entered into cells and thus also behaved as a CPP, it failed to accumulate at nucleoli and was not toxic for U2OS cells (**Fig. S2B,C**). Collectively, the experiments presented above revealed that arginine-rich DPRs have a general effect on cellular reactions that use nucleic acid substrates.

**Fig. 2.**
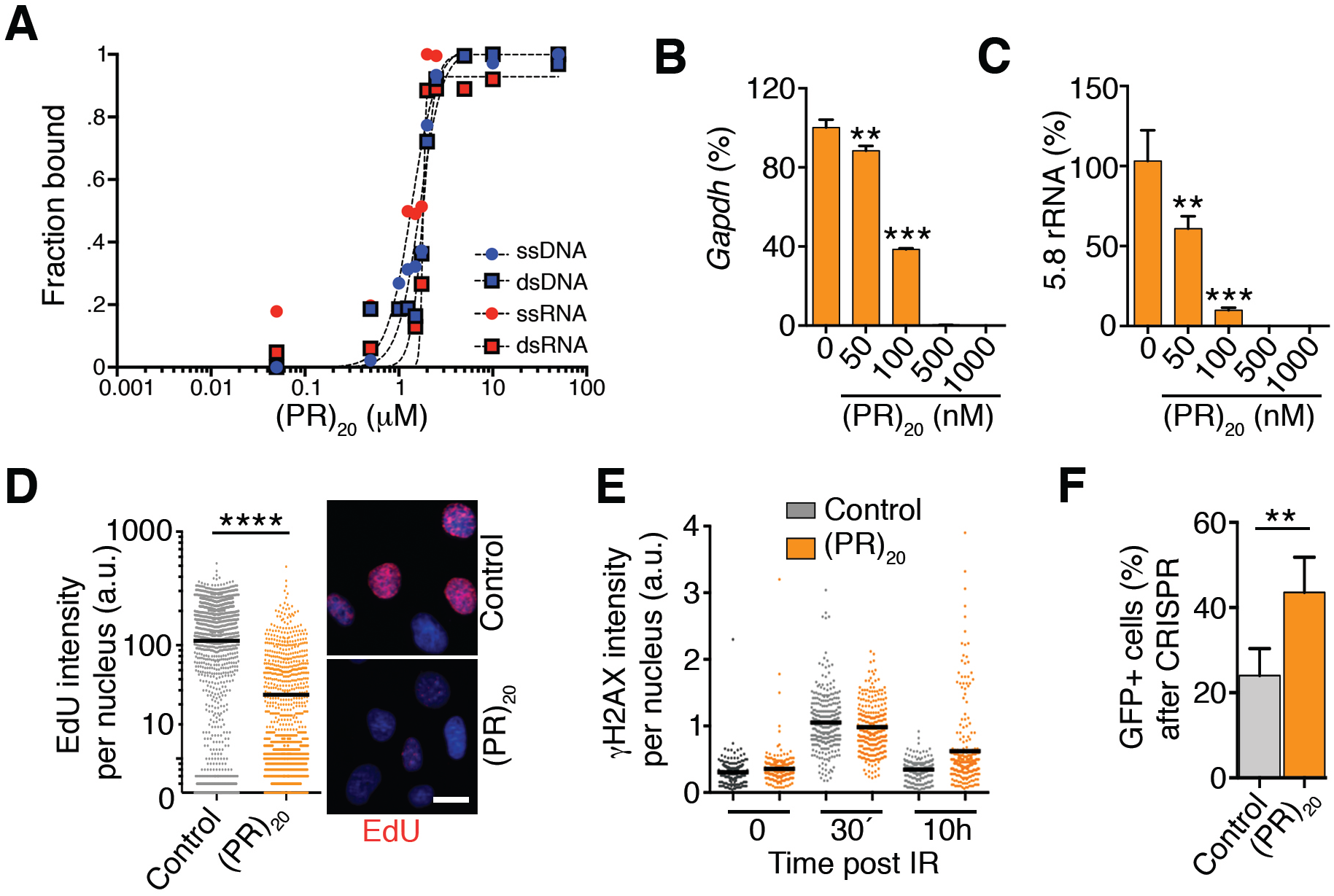
Effects of (PR)_20_ on DNA-based reactions. (**A**) Quantification of EMSA assays evaluating the binding of (PR)_20_ to 19 nt-long ssDNA, ssRNA, dsDNA and dsRNA molecules. Each probe (0.2 µM) was incubated with increasing concentrations of (PR)_20_ for 10’. Curve-fitting was performed using non-linear regression with the Hill equation. (**B, C**) Percentage of *GAPDH* or 5.8 rRNA levels quantified by qPCR in reactions containing increasing doses of (PR)_20_ (n=3). (**D**) HTM-mediated analysis of 5-ethynyl-deoxyuridine (EdU) levels per nucleus found in U2OS cells exposed to 10 µM (PR)_20_ for 4 h. EdU was added 30’ prior to fixation. Representative images from these analyses are provided (right). Scale bar (white) indicates 5 µm. (**E**) HTM-mediated analysis of γH2AX levels per nucleus in U2OS cells exposed to 3Gy of IR in the presence or absence of 10 µM (PR)_20_ for the indicated times. Note the accumulation of cells with persistent DNA damage (γH2AX) 10 h after IR in the presence of the DPR. (**F**) Efficiency of CRISPR-mediated gene deletion quantified in mouse embryonic stem cells stably expressing EGFP, a doxycycline-inducible Cas9 and an EGFP-targeting sgRNA. The percentage of EGFP-positive cells was quantified by flow cytometry 72 h after inducing Cas9 expression with doxycycline in the presence or absence of 10 µM (PR)_20_ (n=3). **, p<0.01; ***, p<0.001; ****, p<0.0001. See also **Fig. S2**.

### Rescue of the effects of (PR)_20_ by nucleic acids

Based on the previous results, we investigated if the effects of the (PR)_20_ peptide could be alleviated by the presence of non-coding oligonucleotides that could scavenge the peptide. In support of this, addition of a 646 nucleotide (nt) RNA lacking ATG rescued the effect of (PR)_20_ on *in vitro* translation reactions in a dose-dependent manner (**Fig. 3A**). Next, we evaluated the effect of adding non-coding oligonucleotides to cells exposed to the DPR. Remarkably, 19 or 38 nt-long ssDNA oligonucleotides rescued the toxicity of (PR)_20_ in U2OS cells both in short-term viability experiments as well as in clonogenic assays (**Fig. 3B,C**). Consistently, the presence of the oligonucleotides rescued the effects of the DPR in reducing translation rates (**Fig. S3A**). Importantly, RNA or DNA oligonucleotides did not prevent the entry of the (PR)_20_ into the nucleus or its accumulation at nucleoli (**Fig. 3D and Fig. S3B**). On the contrary, the presence of the DPR led to the entry of ssDNA and, to a lesser extent, ssRNA, into cells, and their accumulation at nucleoli (**Fig. 3E and Fig. S3C**). In fact, oligoarginines have been used as non-lipidic transfection reagents for decades (Emi et al., 1997), and other arginine-rich CPPs such as HIV-I TAT also facilitate the delivery of nucleic acids or proteins into cells or even animals (Schwarze et al., 1999). In summary, the effects of (PR)_20_ can be alleviated by the presence of oligonucleotides that bind to the DPR thereby limiting the effective concentration of the toxic peptide that can interact with cellular nucleic acids.

**Fig. 3.**
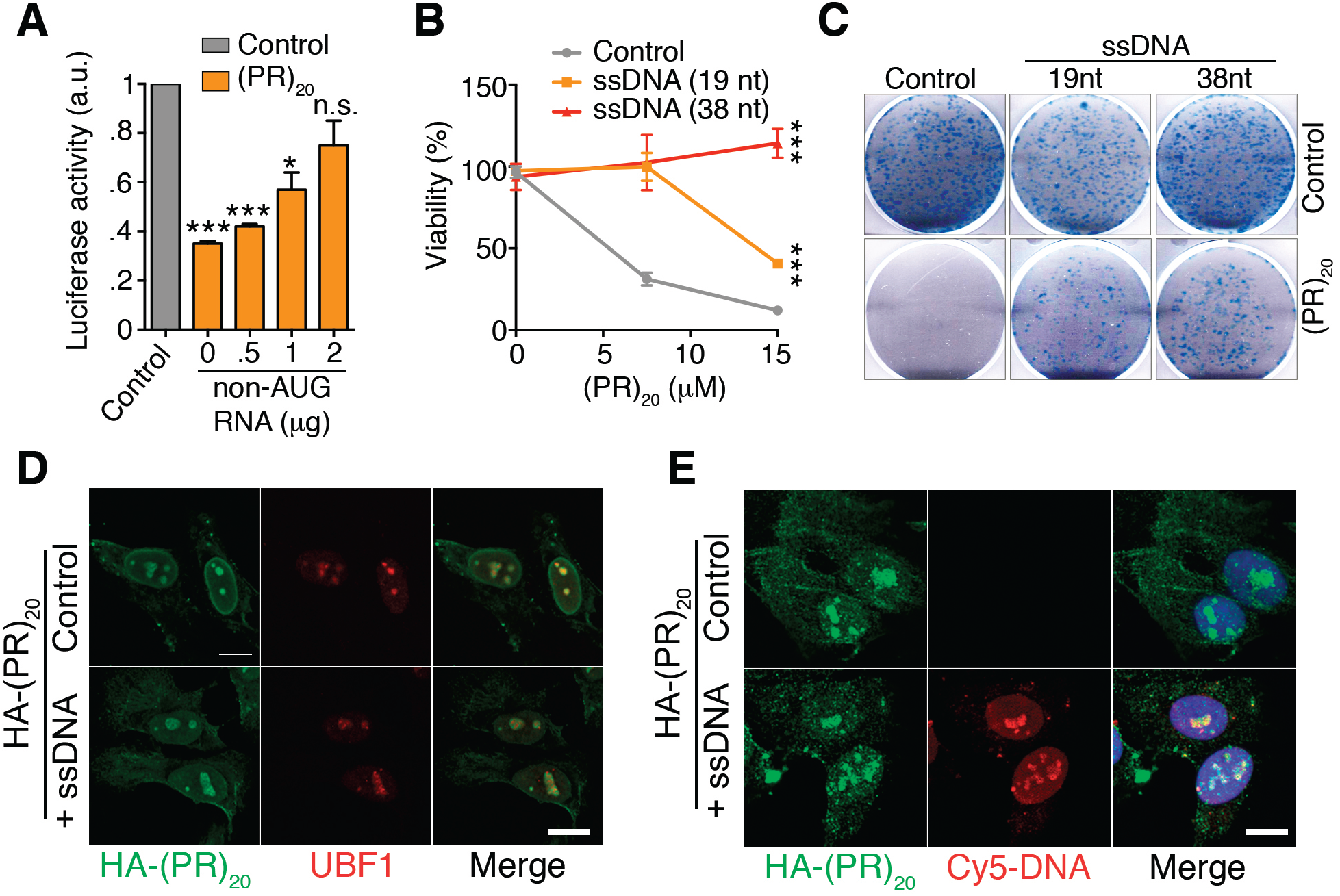
Non-coding nucleic acids rescue the effects of (PR)_20_ DPRs. (**A**) *In* vitro translation of a luciferase mRNA with or without of 0.5 µM (PR)_20_ and in the presence of increasing amounts of a 646 nt non-coding RNA. (**B**) Percentage of viable cells as evaluated with a CellTiter-Glo luminescent assay in U2OS cells treated with increasing doses of (PR)_20_ alone or together with 2 µM of ssDNA oligonucleotides (19 nt or 38 nt) (n=3). (**C**) Clonogenic survival assay of U2OS cells exposed to 7.5 µM (PR)_20_ alone or together with 2 µM of ssDNA oligonucleotides (19 nt or 38 nt). (**D**) Immunofluorescence of HA-(PR)_20_ (green) and the nucleolar factor UBF1 (red) in U2OS cells treated with 7.5 µM HA-(PR)_20_ alone or together with 2 µM of a 19 nt ssDNA oligonucleotide for 8 h. (**E**) Immunofluorescence of HA (green) and Cy5 (red) in U2OS cells treated with 7.5 µM HA-(PR)_20_ alone or together with 4 µM of a Cy5-labeled 19 nt ssDNA oligonucleotide for 8 h. Scale bar (white) in D and E indicates 2.5 µm. *, p<0.1; **, p<0.01; ***, p<0.001. See **Fig. S3** for an HTM-mediated quantification of the image analyses.

### PROTAMINE recapitulates the cellular and biochemical effects of (PR)_20_

We next explored if the effects observed with (PR)_20_ would be similar to those triggered by other arginine-rich CPPs. First, we tested the effects of (GR)_20_, which is the other arginine-rich DPR associated to *C9ORF72* mutations that was reported to behave as a toxic CPP (Kwon et al., 2014). As in the case of (PR)_20_, (GR)_20_ peptides inhibited biochemical reactions using DNA or RNA such as qPCR and RNAse A-mediated RNA degradation (**Fig. S4A,B**). In our hands, however, the effects of (GR)_20_ in cells were more limited likely due to the reported poor stability of this peptide (Kwon et al., 2014). In any case, to provide a more general case on whether the nucleic-acid binding model applies to other arginine-rich CPPs we evaluated the effects of PROTAMINE, a sperm-specific small polypeptide that has the highest percentage of arginine content from the animal proteome (Balhorn, 2007). In addition, PROTAMINE has also been used as a DNA or RNA transfection reagent (Scheicher et al., 2015) and is known to be toxic for poorly understood reasons (Sokolowska et al., 2016). Interestingly, while PROTAMINE is known for its DNA-binding properties during spermatogenesis, EMSA assays revealed a similar affinity of PROTAMINE for DNA and RNA (**Fig. S4C**). In addition, and like (PR)_20_ or (GR)_20_, PROTAMINE had an inhibitory effect on DNA- or RNA-based reactions such as PCR and RNA degradation (**Fig. S4D,E**). In what regards to the cellular effects of PROTAMINE, these were remarkably similar to those observed with (PR)_20_ peptides. Accordingly, PROTAMINE accumulated at nucleoli (**Fig. 4A**) and mimicked the effects triggered by (PR) DPRs in U2OS cells including reduced transcription and translation rates (**Fig. 4B,C**) and alterations in splicing (**Fig. 4D**). In summary, the effects on nucleic acid metabolism observed with ALS-associated (PR)n peptides seem to be a general property of arginine-rich CPPs.

**Fig. 4.**
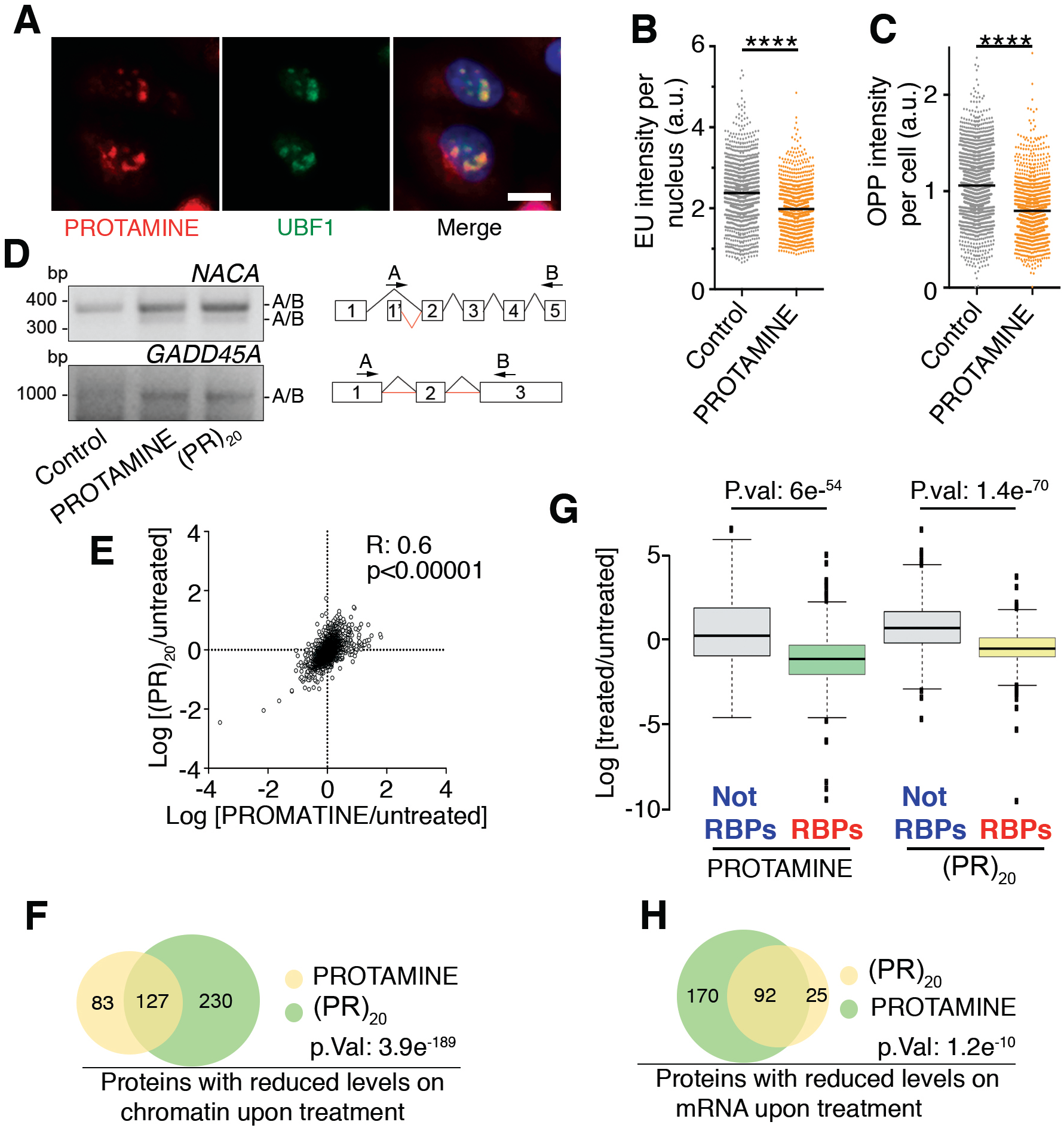
PROTAMINE and (PR)_20_ treatments trigger a widespread displacement of DNA- and RNA-binding factors. (**A**) Immunofluorescence of Cy3-labelled salmon PROTAMINE (red) and the nucleolar factor UBF1 (green) in U2OS cells treated with 30 µM Cy3-PROTAMINE for 4 h. Scale bar (white) indicates 2.5 µm. (**B, C**) HTM-mediated quantification of the EU levels per nucleus (**B**) and O-propargyl-puromycin (OPP) levels per cell (**D**) in U2OS cells exposed to 30 µM PROTAMINE for 16 (**B**) or 24 (**C**) h. EU and OPP were added 30’ prior to fixation. (**D**) RT-PCR of *NACA* and *GADD45A* mRNAs in U2OS cells treated with PROTAMINE (30 µM) or (PR)_20_ (20 µM) using primers in non-consecutive exons to monitor alternative splicing events. A scheme of the primers used is provided on the right side of the panel. See also **Fig. S4A-E** for further analyses or PROTAMINE and a (GR)_20_ peptide. (**E**) Distribution of the chromatin-bound levels of all proteins identified by proteomics in U2OS cells treated with PROTAMINE (30 µM) or (PR)_20_ (20 µM) for 90’. Numbers indicate the Pearson’s correlation coefficient (R) and the p-value for the linear correlation of the changes induced by both treatments. (**F**) Venn diagram illustrating the overlap among the proteins that show reduced levels on chromatin upon treatment of U2OS cells with PROTAMINE (30 µM) or (PR)_20_ (20 µM) for 90’. See also **Table S2**. (**G**) Distribution of the levels of all proteins identified after isolation of the RNA-bound proteome in U2OS cells treated with PROTAMINE (30 µM) or (PR)_20_ (20 µM) for 180’. RNA-binding proteins (RBPs) were identified as those significantly enriched after the isolation of mRNA-binding factors (see also **Fig. S4F,G** for the analysis of this experiment). Note that treatment with either agent leads to a selective reduction in the levels of RBPs bound to mRNA that is not observed on the rest of the factors (Not RBPs). (**H**) Venn diagram illustrating the overlap among the proteins that show reduced levels on mRNA upon treatment of U2OS cells with PROTAMINE (30 µM) or (PR)_20_ (20 µM) for 90’. ****, p<0.0001. See also **Table S2**.

### Exposure to arginine-rich CPPs lead to a generalized displacement of RNA- and DNA-binding factors

Finally, we seek to identify the mechanism by which oligoarginines interfere with cellular reactions using nucleic acids. During spermatogenesis, PROTAMINE binds to DNA and leads to the eviction of histones and High Mobility Group (HMG) proteins from chromatin, allowing a higher compaction of the sperm genome. Given the similar phenotypes triggered by PROTAMINE and arginine-rich CPPs in somatic cells, we reasoned that these could be due to a widespread effect of the peptides in displacing DNA-binding factors from chromatin. Indeed, a proteomic analysis of the changes in chromatin-bound factors that are triggered by PROTAMINE and (PR)_20_ showed a highly statistically significant linear correlation and overlap between both conditions (**Fig. 4E,F**). Moreover, linker histone H1 variants or High Mobility Group (HMG) proteins were among the proteins that showed a highest reduction in their chromatin-bound levels upon (PR)_20_ or PROTAMINE exposure, further supporting that (PR)_20_ peptides phenocopy the effects of PROTAMINE (**Table S2**). Then, and given that PROTAMINE and (PR)_20_ also bind to RNA (**Fig. S4C**), we evaluated if these peptides had a general effect on displacing RNA-binding factors from cellular RNAs. Indeed, through the use of a proteomic pipeline for the analysis of the RNA-bound proteome (Castello et al., 2013), we observed that both PROTAMINE and (PR)_20_ led to an overall reduction in the amount of RNA-binding factors that were bound to RNA (**Fig 4G** and **Fig S4F,G**). In addition, and similar to what happens on chromatin, there was a highly significant correlation between the factors that were displaced from RNA upon treatment with PROTAMINE and (PR)_20_ (**Fig. 4H** and **Table S3**). Noteworthy, the list of proteins that were displaced from RNA included well-known ALS-associated RNA-binding proteins such as TARDBP (TDP43) or FUS. In summary, the presence of arginine-rich CPPs leads to a generalized displacement of RNA- and DNA-binding factors from cellular RNA and chromatin, which provides a general mechanism to explain the widespread defects in nucleic acid metabolism that are observed upon exposure to these peptides.

## DISCUSSION

Facilitating the delivery of nucleic acids or proteins into cells is an intense area of biomedical research, particularly in these days where the use of biological agents is gaining momentum for the treatment of human disease (Bruce and McNaughton, 2017). In this context, while arginine-rich oligopeptides are among the most efficient CPPs, they also confront with toxicity as has been observed during the medical use of PROTAMINE. The discovery of toxic arginine-rich CPPs in a subset of ALS/FTD patients has now made the study of this phenomenon even more relevant. The work presented here provides a unifying mechanism to explain the toxic effects of arginine-rich CPPs, including those translated from ALS-associated *C9ORF72* intronic repeat expansions. We propose that the high affinity of oligoarginines for RNA or DNA leads to a widespread coating of nucleic acids in cells, with a consequent generalized displacement of RNA- or DNA-binding factors from chromatin and RNA. This model implies that any reaction using nucleic acid substrates including RNA transcription, splicing and translation or DNA replication and repair will be affected by arginine-rich CPPs, and in fact effects on all of these cellular reactions have been reported in ALS/FTD (Hardiman et al., 2017). Interestingly, recent work showed that a GFP-(PR)_50_ peptide accumulated at heterochromatin leading to the loss of HP1*α* (Zhang et al., 2019). While this effect was proposed to be due to a specific interaction of poly(PR) peptides with HP1*α* that disrupted its liquid-liquid phase separation properties, we think it could be explained by a more general effect of oligoarginines on displacing chromatin-bound factors. In the light of our observations, we propose that the cellular pathology driven by the presence of (PR)n and (GR)n DPRs in *C9ORF72* mutant patients, would be mechanistically equivalent to what could be expected if PROTAMINE were to be accidentally expressed in adult cells.

In what regards as to how the widespread effects of arginine-rich CPPs could be limited, we here show that these can be alleviated by non-coding oligonucleotides that, through binding to the peptides, can limit the effective concentration available to interact with cellular nucleic acids. While the efficient delivery of non-coding oligonucleotides into neurons seems challenging, it should be noted that recent treatments for other neuromuscular disorders such as spinal muscular atrophy are based on the direct injection of modified non-coding antisense oligonucleotides into the cerebrospinal fluid (Wood et al., 2017). Besides oligonucleotides, some other hints as to how to limit the toxicity of arginine-rich CPPs can be obtained from our knowledge on how these peptides enter into cells. Even though a full understanding is still missing, several works have shown that the internalization of arginine-rich CPPs occurs through binding to heparanated receptors on the cell membrane (Fuchs and Raines, 2006). In addition, and as mentioned, PROTAMINE is medically used as an antidote of the anticoagulant heparin. In this context, heparin or derivatives could be explored as a strategy to limit the toxicity of arginine-rich CPPs. While heparin delivery into cells might not be trivial, these strategies might be particularly relevant in the context of paracrine effects, which have been reported to be relevant not only in *C9ORF72* derived DPRs (Westergard et al., 2016; Zhou et al., 2017) but also for AIDS-related neurodegeneration driven by the TAT protein from the HIV-1 virus (King et al., 2006). In summary, the work presented here provides a unifying mechanism to explain the widespread effects of arginine-rich CPPs in nucleic acid metabolism, and provides some initial proof-of-principle ideas as to how this knowledge could be used to limit their toxicity in mammalian cells.

## ACKNOWLEDGEMENTS

We would want to thank Drs. André Nussenzweig an Jordi Carreras-Puigvert for insightful comments on the manuscript. Research was funded by Fundación Botín, by Banco Santander through its Santander Universities Global Division and by grants from the Spanish Ministry of Economy and Competitiveness (MINECO) (SAF2014-54498-R, co-financed with European FEDER funds), Spanish Ministry of Science, Innovation and Universities (RTI2018-102204-B-I00, co-financed with European FEDER funds) and the European Research Council (ERC-617840) to OF; DKFZ NCT3.0 Integrative Project in Cancer Research grant (NCT3.0_2015.54 DysregPT) and SFB 1036/TP07 from the Deutsche Forschungsgemeinschaft to G.S.

## AUTHOR CONTRIBUTIONS

V.L. and O.S. contributed to most experiments and data analyses and to the preparation of the figures. I.D. and I.V. helped with in vitro translation and viral infection experiments. E.Z. and J.M. helped with proteomic analyses. M. H. and G. S. conducted polysome analyses and B.J. generated RPS9^SBP^-expressing HeLa cells. J.B. performed electron microscopy experiments. R.F. helped with DNA and RNA binding experiments. O.F. supervised the study and wrote the MS.

## DECLARATION OF INTERESTS

The authors declare no competing interests.

## MATERIALS AND METHODS

### Cell culture

U2OS, BHK-21, RPE, HeLa-SBP and HeLa-RPS9^SBP^ cells stably expressing RPS9-Flag-TEV-SBP were cultivated in standard DMEM medium supplemented with 10% FBS, 2 mM L-glutamine and 1% penicillin/streptomycin. The previously described mouse ES^Cas9^ cells (Ruiz et al., 2016) were grown on gelatin in DMEM (high glucose) supplemented with 15% knockout serum replacement (Invitrogen), LIF (1000 U/ml), 0.1 mM non-essential amino acids, 1% glutamax and 55 mM β-mercaptoethanol.

### Treatments with peptides or oligonucleotides

Peptides containing 20 PR dipeptide repeats and a C-terminal HA epitope tag were synthesized at Genscript. Hydroxyurea (H8627) and salmon PROTAMINE (P4005) were obtained from Sigma and cycloheximide (239764) from Calbiochem. Fluorophore-labelled oligonucleotides were synthesized by Sigma, with the following sequences: Cy3-5’-DNA (CCACTGCACCGCTGCTAGG); Cy5-5’-DNA (CCTAGCAGCGGTTGCAGTGG); Cy3-5’-RNA (CCACUGCACCGCUGCUAGG) and Cy5-RNA (CCUAGCAGCGGUUGCAGUGG).

### Immunofluorescence and High Throughput Microscopy

For immunofluorescence using HA, coilin and UBF antibodies, cells were fixed with 4% PFA prepared in PHEM buffer (60 mM Pipes, 25 mM Hepes, 10 mM EGTA, 2 mM MgCl_2_ pH 6.9) containing 0.2% of Triton X-100, and permeabilized with 0.5% Triton X-100 after fixation. For HTM, cells were grown on *µ*CLEAR bottom 96-well plates (Greiner Bio-One) and immunofluorescence of γH2AX was performed using standard procedures. Analysis of DNA replication by EdU, transcription by EU and translation by OPP incorporation were done using Click-It kits (Invitrogen) following manufacturer’s instructions. In all cases, images were automatically acquired from each well using an Opera High-Content Screening System (Perkin Elmer). A 20x or 40x magnification lens was used and images were taken at non-saturating conditions. Images were segmented using DAPI signals to generate masks matching cell nuclei from which the mean signals for the rest of the stainings were calculated.

### Cell viability

4,000 cells were seeded per well in a 96-well tissue culture plate and treated with the indicated concentrations of (PR)_20_ alone or together with 4 µM of 19 or 38 nt-long Cy5-RNA or Cy5-DNA oligonucleotides. 36 hours after the treatment, cell viability was measured using a luminescent system (CellTiter-Glo, Promega), according to the manufacturer’s protocol. Viability is plotted as percentages compared to untreated controls.

### Clonogenic assays

2,000 U2OS cells were plated in 6-well tissue culture plates in culture medium. The following day, cells were incubated with 10 μM (PR)_20_ alone or in combination with 2 µM of 19 (CCACTGCACCGCTGCTAGG) or 38 (CCACTGCACCGCTGCTAGGATCGCCTGAAATCGTTGGC) nucleotide-length ssDNA molecules. After 2 days the medium was changed and cells were then grown for 8 additional days in untreated medium. At the end of the experiments, cells were fixed and stained with 0.4% methylene blue in methanol for 30 min.

### Peptide labelling

The Cy^®^3 Mono 5-pack kit (Sigma) was used for the labeling of PROTAMINE. Briefly, 0.150 mM salmon PROTAMINE was incubated with 0.850 mM of Cy3 NHS Ester in 20 mM Hepes and 150 mM NaCl buffer overnight at 4°C, followed by the addition of 1 M Tris pH7.5 in order to quench the reaction. Cy3-protamine was purified using 3K Amicon Ultra 0.5 mL centrifugal filters (Sigma).

### Immunoprecipitation

10×10^6^ Hela-SBP and Hela-RPS9^SBP^ cells were lysed in cold RNA-IP lysis buffer (50 mM Tris pH 8.0, 150 mM NaCl, 1 mM MgCl_2_, 2 % NP-40, 10% glycerol and freshly added Complete protease inhibitors) and incubated with 30 μL of Strep-Tactin beads (IBA) for 1 h at 4°C with constant shaking. Beads were washed 6 times in NET2 buffer (50 mM Tris pH 7.5, 150 mM NaCl, 0.5% Triton X-100).

### Chromatin fractionation

After treatment with PR20 or PROTAMINE, cells were washed twice with ice-cold phosphate-buffered saline (PBS), resuspended in 180 μl of ice-cold hypotonic lysis buffer (10 mM HEPES, pH 7.9, 10 mM KCl, 0.1 mM EDTA containing protease and phosphatase inhibitors), and incubated on ice for 10 min, followed by addition of 20 μl of Nonidet P-40; after 3 min at room temperature, cells were vortexed and the cytosolic fraction was obtained by centrifugation for 5 min at 2,500 × g. The nuclear pellet was resuspended in high-salt-concentration extraction buffer (20 mM HEPES, pH 7.9, 0.4 M NaCl, 1 mM EDTA containing protease and phosphatase inhibitors) and incubated with shaking at 4°C for 1 h. For chromatin fraction the pellet was then extracted in 50 mM Tris, pH 7.5, 8 M urea, and 1% CHAPS.

### Isolation of RNA-binding proteins

The systems-wide purification of RNA-binding proteins was developed as performed reported (Castello et al., 2013). Briefly, 5×150 cm2 dishes of 60 % confluent U2OS cells were treated for 4 hours with 20 *υ*M of PR_20_ or Protamine. RBPs and polyadenylated RNAs were crosslinked by irradiating the cells with 0.15 J/cm^2^ (∼1 min) at 254 nm UV, and non-irradiated cells were used as a control. Cells were lysated and proteins covalently to mRNA were captured with oligo(dT) magnetic beads. After stringent washes, the mRNA interactome was determined by quantitative mass spectrometry (MS).

### Mass spectrometry

IP samples were eluted in urea, FASP-digested with Lys-C/trypsin and analyzed by LC-MS/MS. Label-free quantification was performed using MaxQuant. Chromatin and whole cell extract samples were trypsin-digested using S-traps, isobaric-labelled with iTRAQ 8-plex (chromatin) or TMT 11-plex (whole cell) reagents and pre-fractionated with RP-HPLC at high pH. Fractions were then analyzed by LC-MS/MS and raw data was processed with MaxQuant. Statistical analyses were performed with Perseus and ProStar.

For RNA-bound proteomics proteins were digested by means of standard FASP protocol. Briefly, proteins were reduced (15 mM TCEP, 30 min, RT), alkylated (50 mM CAA, 20 min in the dark, RT) and sequentially digested with Lys-C (Wako) (o/n at RT) and trypsin (Promega) (6 h at 37 °C). Resulting peptides were desalted using Sep-Pak C18 cartridges (Waters). LC-MS/MS was done by coupling an UltiMate 3000 HPLC system to a Q Exactive Plus mass spectrometer (Thermo Fisher Scientific). Peptides were loaded into a trap column Acclaim™ PepMap™ 100 C18 LC Columns 5 *µ*m, 20 mm length) for 3 min at a flow rate of 10 *µ*l/min in 0.1% FA. Then peptides were transferred to an analytical column (PepMap RSLC C18 2 *µ*m, 75 *µ*m × 50 cm) and separated using a 89 min effective curved gradient (buffer A: 0.1% FA; buffer B: 100% ACN, 0.1% FA) at a flow rate of 250 nL/min from 2% to 42.5% of buffer B. The mass spectrometer was operated in a data-dependent mode, with an automatic switch between MS (350-1400 m/z) and MS/MS scans using a top 15 method (intensity threshold signal ≥ 3.8e4, z ≥2). An active exclusion of 26.3 s was used. Peptides were isolated using a 2 Th window and fragmented using higher-energy collisional dissociation (HCD) with a normalized collision energy of 27. Protein interaction networks arising from proteomic experiments were analyses with STRING (Szklarczyk et al., 2015).

Mass spectrometry proteomics data associated to this work have been deposited to the ProteomeXchange Consortium via the PRIDE (Vizcaino et al., 2016) partner repository with the dataset identifier PXD010555 (Username; Password: ZnXyUNzE) and PXD014085 (Username; Password: UAy57afS).

### Polysome analyses

Hela-RPS9^SBP^ cells were treated with either water as control or 10 *µ*M of (PR)_20_ during 16 hours. Ribosomes were stalled by addition of 100 *µ*g/mL cycloheximide (CHX) for 5 min, and cells were lysed in polysome lysis buffer (15 mM Tris-HCl pH 7.4, 15 mM MgCl_2_, 300 mM NaCl, 1% Triton-X-100, 0.1% β-mercaptoethanol, 200 U/mL RNAsin (Promega), 1 complete Mini Protease Inhibitor Tablet (Roche) per 10 mL). Nuclei were removed by centrifugation (9300×G, 4°C, 10 min) and the cytoplasmic lysate was loaded onto a sucrose density gradient (17.5–50% in 15 mM Tris-HCl pH 7.4, 15 mM MgCl_2_, 300 mM NaCl and, for fractionation from BMDM, 200 U/ml Recombinant RNAsin Ribonuclease Inhibitor, Promega). After ultracentrifugation (2.5 h, 35,000 rpm at 4°C in a SW60Ti rotor), gradients were eluted with a Teledyne Isco Foxy Jr. system into 16 fractions of similar volume.

### In vitro ribosome assembly

Ribosomal subunits were purified from BHK21 cells using previously described procedures (Pisarev et al., 2007). Briefly, the ribosomal fraction (P100) was treated with 1 mM puromycin for 15 min on ice and then for 15 min at 37°C. The suspension was adjusted to 0.5 M KCl and loaded onto 10 to 30% sucrose density gradients which were centrifuged at 22,000 rpm for 16 h at 4°C in a SW40 rotor. The peaks corresponding to 40S and 60S subunits were identified by optical density at 260 nm. Finally, fractions were concentrated using YM-100 centricons. 1 pmol of 40S and 60S were incubated in assembly buffer (25 mM of TrisHCl pH7.5, 100 mM KCl, 5 mM MgCl_2_ and 3mM of DTT) for 45 minutes at 30 °C. To control for non-assembly, the concentration of MgCl_2_ in the buffer was reduced to 1 mM.

### Electron microscopy

For negative staining 3 µl of purified 40S+60S ribosome fractions or alternatively 3 µl of 40S+60S ribosome fractions in the presence of (PR)_20_ (1:10 molar ratio) were deposited onto freshly glow-discharged carbon-coated 400 mesh copper electron microscopy (EM) grids (Electron Microscopy Sciences) and retained on the grid for 1 min. Afterwards, grids were washed with three distinct 50 μL drops of MilliQ water. The excess of water was removed with filter paper and the grids were placed on the top of three different 40 μL drops of 2% uranyl acetate and stained on the last drop for 1 min. The grids were then gently stripped for 4 s and air dried. Finally, grids were visualized on a Tecnai 12 transmission electron microscope (Thermo Fisher Scientific Inc.) with a lanthanum hexaboride cathode operated at 120 keV. Images were recorded at 61320 nominal magnification on a 4k×4k TemCam-F416 CMOS camera (TVIPS).

### In vitro translation

In vitro translation was performed in nuclease-treated rabbit reticulocyte lysates (RRL) (Promega cat #L4960) using a luciferase mRNAs in the presence of the indicated concentrations of (PR)_20_. Reactions were incubated for 60 mins at 30°C and a 5 µl alicuot was used to measure luciferase activity in 100 µl of reaction buffer (15mM K2HPO4, 15mM MgSO4, 4mM EGTA pH 8, 4mM DTT, 2mM ATP and 0.1mM luciferin) in a Berthold Luminometer. For radioactive measurements, in vitro translation was performed in nuclease-treated RRL (Promega) using 150 ng of luciferase mRNA in the presence of 15 μCi of [35S]-Met for 60 min at 30°C. Samples were denatured in sample buffer and analyzed by SDS-PAGE and autoradiography.

### In situ hybridization

In situ hybridization was carried out as previously described (Palanca et al., 2014). Briefly, U2OS cells were fixed with 4% PFA prepared in PHEM buffer. An oligo dT_(50)-_mer, 5′-end labeled with biotin (MWG-Biotech, Germany) was used as a probe for in situ hybridization to poly(A) RNA. After hybridization, cells were washed and the hybridization signal was detected with FITC-avidin for 30 min. All samples were mounted with the ProLong anti-fading medium (Invitrogen).

### RNA degradation assay

RNA was extracted from mouse embryonic stem cells using the Absolutely RNA microprep kit (Agilent). 1 *µ*g of RNA was pre-incubated with 4.5 *µ*g of (PR)_20_ or water for 10 min at room temperature, and subsequently treated with serially diluted RNase A (Qiagen) at 37°C for 15 min, at the end of which the reaction was halted by RNasin ribonuclaease inhibitor (Promega) and 1% SDS. The resulting mixture was column-purified using the RNA Clean & Concentrator-5 kit (Zymo Research), according to manufacturer’s instructions. The recovered RNA was analyzed by agarose gel electrophoresis, and quantified by image analysis (ImageJ, NIH, USA).

### Reverse transcription

500 ng of RNA isolated from mESC was pre-incubated on ice for 5 minutes with increasing concentrations of (PR)_20_. The mixture was then used as template for cDNA synthesis using the SuperScript III First Strand Synthesis System (ThermoFisher Scientific) according to manufacturer’s instructions. The reaction mixture was subsequently treated with RNase A (15 min at 37°C), solubilized with SDS 1%, and finally purified using PCR cleanup columns (Qiagen). The yield of cDNA was visualized by agarose gel electrophoresis, and quantified by fluorometry using the Qubit kit (ThermoFisher Scientific), according to manufacturer’s instructions.

### RT-PCR

cDNA generated from mESC was pre-incubated on ice for 5 min with increasing concentrations of (PR)_20_. Out of this mixture, 50 ng of cDNA were used in each reaction. RT-PCR was carried out using the 2X SYBR Select Master Mix (Life Technologies) in MicroAmp© Optical 384-Well plates (Applied Biosystems) in the QuantStudio™ 6 Flex Real-Time PCR System (Thermo Fisher) using standard protocols. The sequences of the primers used are as follows: *Gapdh*-F (TTCACCACCATGGAGAAGGC), *Gapdh*-R CCCTTTTGGCTCCACCCT, 5.8S-F (GTCGATGAAGAACGCAGCTA) and 5.8S-R (AACCGACGCTCAGACAGG).

### Viral infection

BHK-21 cells were infected with Sindbis virus (Alphavirus, ssRNA genome with positive polarity) at a MOI of 20 viral PFU/cell in the absence or presence of (PR)_20_. Cells were collected 4 hours after infection and total RNA was purified using RNeasy Mini Kit (Quiagen). Virus-specific primers (F:GGTACTGGAGACGGATATCGC and R:CGATCAAGTCGAGTAGTGGTTG) were used to detect viral RNA by qRT-PCR, which were normalized against cellular *GAPDH*.

### Splicing

U2OS cells were plated at a density of 2.4 × 10^4^ cells/mL on 6-well plates and treated on the following day with either 20 μM of (PR)_20_ or 30 *µ*M protamine. RT-PCR was performed as described above using previously described primers to detect alternative splicing events at *GADD45A* and *NACA* mRNAs (Kwon et al., 2014).

### EMSA

To analyze the binding of (PR)_20_ to ssDNA and ssRNA, 0.2 µM of Cy3-RNA (CCACUGCACCGCUGCUAGG) or Cy3-DNA (CCACTGCACCGCTGCTAGG) oligonucleotides were incubated with increasing amounts of (PR)_20_ for 10 min at room temperature. To evaluate the binding of (PR)_20_ to dsDNA and dsRNA, Cy3-RNA and Cy3-DNA oligonucleotides were annealed with the complementary RNA (CCUAGCAGCGGUUGCAGUGG) or DNA (CCTAGCAGCGGTTGCAGTGG) molecule prior to incubation with (PR)_20_. Reactions were supplemented with 4 μL of 6x loading buffer (30% glycerol, bromphenol blue, xylene cyanol) and were resolved by polyacrylamide gel electrophoresis in 4-12% TBE Gels (Invitrogen) in 1%TBE buffer at 100 V for 45 min at 4°C. The gels were scanned and Cy3 intensity was measured using Typhoon Trio (GE Health Care). Band quantification was performed using ImageJ (Schneider et al., 2012) and data fitting was performed using Graphpad Prism 7.

### CRISPR/Cas9 efficiency

The previously described ES^Cas9^ mESC line (Ruiz et al., 2016) which carries a Doxycycline-inducible Cas9 was co-infected with lentiviral vectors expressing EGFP (pLVTHM, Addgene, 12247) and an EGFP-targeting sgRNA cloned in the pKLV-U6gRNA-PGKpuro2ABFP backbone (Addgene, 50946). Two independent clones with stable expression of all components were seeded on gelatin. After six hours, Doxycycline (1 *µ*g/mL) and/or (PR)_20_ (10 *µ*M) were added to the medium. 72 hours later, cells were recovered and the percentage of GFP-positive cells was quantified by flow cytometry using the FlowJo software (BD).

### Phosphatase assay

Briefly, RPE cells were lysate with RIPA (150 mM NaCl, 25 mM Tris-Hcel, 1 mM EDTA, 0.5 mM EGTA, 1% Triton X-100, 0.1 % SDS and protease inhibitors) and 100 mg of protein was used for each condition. PP2A Immunoprecipitation Phosphatase Assay kit was used for determining phosphatase activity following manufactured instructions. The dephosphorylation of a KRpTIRR peptide activity was measured by absorbance of Malakita green.

## Notes

#### Summary of Updates

In the new version, we have included additional proteomic data to reveal that arginine-rich peptides, including PROTAMINE or the ALS-related poly(PR) peptides lead to a generalized displatement of RNA-binding factors from mRNA. In addition, we have included additional data to show the effect of other arginine rich peptides (poly(GR), show that these effects are not shared by lysin-rich CPPs (such as poly(PK) and illustrate that these effects of arginine-rich CPPs are restricted to reactions that use nucleic acid templates. Finally, we have changed the overall structure of the manuscript to further indicate that our manuscript is not only related to ALS, but rather provides a general model to explain why arginine-rich cell-penetrating peptides are toxic for animal cells.

## REFERENCES

Ash, P.E., Bieniek, K.F., Gendron, T.F., Caulfield, T., Lin, W.L., Dejesus-Hernandez, M., van Blitterswijk, M.M., Jansen-West, K., Paul, J.W., 3rd, Rademakers, R., et al. (2013). Unconventional translation of C9ORF72 GGGGCC expansion generates insoluble polypeptides specific to c9FTD/ALS. Neuron 77, 639–646.

Balhorn, R. (2007). The protamine family of sperm nuclear proteins. Genome Biol 8, 227.

Boeynaems, S., Bogaert, E., Kovacs, D., Konijnenberg, A., Timmerman, E., Volkov, A., Guharoy, M., De Decker, M., Jaspers, T., Ryan, V.H., et al. (2017). Phase Separation of C9orf72 Dipeptide Repeats Perturbs Stress Granule Dynamics. Mol Cell 65, 1044–1055 e1045.

Borrelli, A., Tornesello, A.L., Tornesello, M.L., and Buonaguro, F.M. (2018). Cell Penetrating Peptides as Molecular Carriers for Anti-Cancer Agents. Molecules 23.

Bruce, V.J., and McNaughton, B.R. (2017). Inside Job: Methods for Delivering Proteins to the Interior of Mammalian Cells. Cell Chem Biol 24, 924–934.

Castello, A., Horos, R., Strein, C., Fischer, B., Eichelbaum, K., Steinmetz, L.M., Krijgsveld, J., and Hentze, M.W. (2013). System-wide identification of RNA-binding proteins by interactome capture. Nat Protoc 8, 491–500.

Chai, N., and Gitler, A.D. (2018). Yeast screen for modifiers of C9orf72 poly(glycine-arginine) dipeptide repeat toxicity. FEMS Yeast Res 18.

Chew, J., Gendron, T.F., Prudencio, M., Sasaguri, H., Zhang, Y.J., Castanedes-Casey, M., Lee, C.W., Jansen-West, K., Kurti, A., Murray, M.E., et al. (2015). Neurodegeneration. C9ORF72 repeat expansions in mice cause TDP-43 pathology, neuronal loss, and behavioral deficits. Science 348, 1151–1154.

DeJesus-Hernandez, M., Mackenzie, I.R., Boeve, B.F., Boxer, A.L., Baker, M., Rutherford, N.J., Nicholson, A.M., Finch, N.A., Flynn, H., Adamson, J., et al. (2011). Expanded GGGGCC hexanucleotide repeat in noncoding region of C9ORF72 causes chromosome 9p-linked FTD and ALS. Neuron 72, 245–256.

DeRouchey, J., Hoover, B., and Rau, D.C. (2013). A comparison of DNA compaction by arginine and lysine peptides: a physical basis for arginine rich protamines. Biochemistry 52, 3000-3009.

Emi, N., Kidoaki, S., Yoshikawa, K., and Saito, H. (1997). Gene transfer mediated by polyarginine requires a formation of big carrier-complex of DNA aggregate. Biochem Biophys Res Commun 231, 421–424.

Frankel, A.D., and Pabo, C.O. (1988). Cellular uptake of the tat protein from human immunodeficiency virus. Cell 55, 1189–1193.

Fuchs, S.M., and Raines, R.T. (2006). Internalization of cationic peptides: the road less (or more?) traveled. Cell Mol Life Sci 63, 1819–1822.

Green, M., and Loewenstein, P.M. (1988). Autonomous functional domains of chemically synthesized human immunodeficiency virus tat trans-activator protein. Cell 55, 1179–1188.

Hardiman, O., Al-Chalabi, A., Chio, A., Corr, E.M., Logroscino, G., Robberecht, W., Shaw, P.J., Simmons, Z., and van den Berg, L.H. (2017). Amyotrophic lateral sclerosis. Nat Rev Dis Primers 3, 17085.

Kanekura, K., Yagi, T., Cammack, A.J., Mahadevan, J., Kuroda, M., Harms, M.B., Miller, T.M., and Urano, F. (2016). Poly-dipeptides encoded by the C9ORF72 repeats block global protein translation. Hum Mol Genet 25, 1803–1813.

King, J.E., Eugenin, E.A., Buckner, C.M., and Berman, J.W. (2006). HIV tat and neurotoxicity. Microbes Infect 8, 1347–1357.

Kwon, I., Xiang, S., Kato, M., Wu, L., Theodoropoulos, P., Wang, T., Kim, J., Yun, J., Xie, Y., and McKnight, S.L. (2014). Poly-dipeptides encoded by the C9orf72 repeats bind nucleoli, impede RNA biogenesis, and kill cells. Science 345, 1139–1145.

Lin, Y., Mori, E., Kato, M., Xiang, S., Wu, L., Kwon, I., and McKnight, S.L. (2016). Toxic PR Poly-Dipeptides Encoded by the C9orf72 Repeat Expansion Target LC Domain Polymers. Cell 167, 789–802 e712.

Lopez-Gonzalez, R., Lu, Y., Gendron, T.F., Karydas, A., Tran, H., Yang, D., Petrucelli, L., Miller, B.L., Almeida, S., and Gao, F.B. (2016). Poly(GR) in C9ORF72-Related ALS/FTD Compromises Mitochondrial Function and Increases Oxidative Stress and DNA Damage in iPSC-Derived Motor Neurons. Neuron 92, 383–391.

Mascotti, D.P., and Lohman, T.M. (1997). Thermodynamics of oligoarginines binding to RNA and DNA. Biochemistry 36, 7272–7279.

Mizielinska, S., Gronke, S., Niccoli, T., Ridler, C.E., Clayton, E.L., Devoy, A., Moens, T., Norona, F.E., Woollacott, I.O.C., Pietrzyk, J., et al. (2014). C9orf72 repeat expansions cause neurodegeneration in Drosophila through arginine-rich proteins. Science 345, 1192–1194.

Mori, K., Weng, S.M., Arzberger, T., May, S., Rentzsch, K., Kremmer, E., Schmid, B., Kretzschmar, H.A., Cruts, M., Van Broeckhoven, C., et al. (2013). The C9orf72 GGGGCC repeat is translated into aggregating dipeptide-repeat proteins in FTLD/ALS. Science 339, 1335–1338.

Nizami, Z., Deryusheva, S., and Gall, J.G. (2010). The Cajal body and histone locus body. Cold Spring Harb Perspect Biol 2, a000653.

Palanca, A., Casafont, I., Berciano, M.T., and Lafarga, M. (2014). Reactive nucleolar and Cajal body responses to proteasome inhibition in sensory ganglion neurons. Biochim Biophys Acta 1842, 848–859.

Park, J., Ryu, J., Kim, K.A., Lee, H.J., Bahn, J.H., Han, K., Choi, E.Y., Lee, K.S., Kwon, H.Y., and Choi, S.Y. (2002). Mutational analysis of a human immunodeficiency virus type 1 Tat protein transduction domain which is required for delivery of an exogenous protein into mammalian cells. J Gen Virol 83, 1173–1181.

Pisarev, A.V., Unbehaun, A., Hellen, C.U., and Pestova, T.V. (2007). Assembly and analysis of eukaryotic translation initiation complexes. Methods Enzymol 430, 147–177.

Renton, A.E., Majounie, E., Waite, A., Simon-Sanchez, J., Rollinson, S., Gibbs, J.R., Schymick, J.C., Laaksovirta, H., van Swieten, J.C., Myllykangas, L., et al. (2011). A hexanucleotide repeat expansion in C9ORF72 is the cause of chromosome 9p21-linked ALS-FTD. Neuron 72, 257–268.

Rossi, S., Serrano, A., Gerbino, V., Giorgi, A., Di Francesco, L., Nencini, M., Bozzo, F., Schinina, M.E., Bagni, C., Cestra, G., et al. (2015). Nuclear accumulation of mRNAs underlies G4C2-repeat-induced translational repression in a cellular model of C9orf72 ALS. J Cell Sci 128, 1787–1799.

Ruiz, S., Mayor-Ruiz, C., Lafarga, V., Murga, M., Vega-Sendino, M., Ortega, S., and Fernandez-Capetillo, O. (2016). A Genome-wide CRISPR Screen Identifies CDC25A as a Determinant of Sensitivity to ATR Inhibitors. Mol Cell 62, 307–313.

Ryser, H.J., and Hancock, R. (1965). Histones and basic polyamino acids stimulate the uptake of albumin by tumor cells in culture. Science 150, 501–503.

Scheicher, B., Schachner-Nedherer, A.L., and Zimmer, A. (2015). Protamine-oligonucleotide-nanoparticles: Recent advances in drug delivery and drug targeting. Eur J Pharm Sci 75, 54–59.

Schneider, C.A., Rasband, W.S., and Eliceiri, K.W. (2012). NIH Image to ImageJ: 25 years of image analysis. Nat Methods 9, 671–675.

Schwarze, S.R., Ho, A., Vocero-Akbani, A., and Dowdy, S.F. (1999). In vivo protein transduction: delivery of a biologically active protein into the mouse. Science 285, 1569–1572.

Smull, C.E., and Ludwig, E.H. (1962). Enhancement of the plaque-forming capacity of poliovirus ribonucleic acid with basic proteins. J Bacteriol 84, 1035–1040.

Sokolowska, E., Kalaska, B., Miklosz, J., and Mogielnicki, A. (2016). The toxicology of heparin reversal with protamine: past, present and future. Expert Opin Drug Metab Toxicol 12, 897–909.

Sprunt, K., Redman, W.M., and Alexander, H.E. (1959). Infectious ribonucleic acid derived from enteroviruses. Proc Soc Exp Biol Med 101, 604–608.

Stopford, M.J., Higginbottom, A., Hautbergue, G.M., Cooper-Knock, J., Mulcahy, P.J., De Vos, K.J., Renton, A.E., Pliner, H., Calvo, A., Chio, A., et al. (2017). C9ORF72 hexanucleotide repeat exerts toxicity in a stable, inducible motor neuronal cell model, which is rescued by partial depletion of Pten. Hum Mol Genet 26, 1133–1145.

Swaminathan, A., Bouffard, M., Liao, M., Ryan, S., Callister, J.B., Pickering-Brown, S.M., Armstrong, G.A.B., and Drapeau, P. (2018). Expression of C9orf72-related dipeptides impairs motor function in a vertebrate model. Hum Mol Genet 27, 1754–1762.

Szklarczyk, D., Franceschini, A., Wyder, S., Forslund, K., Heller, D., Huerta-Cepas, J., Simonovic, M., Roth, A., Santos, A., Tsafou, K.P., et al. (2015). STRING v10: protein-protein interaction networks, integrated over the tree of life. Nucleic Acids Res 43, D447–452.

Tan, R., and Frankel, A.D. (1995). Structural variety of arginine-rich RNA-binding peptides. Proc Natl Acad Sci U S A 92, 5282–5286.

Vizcaino, J.A., Csordas, A., del-Toro, N., Dianes, J.A., Griss, J., Lavidas, I., Mayer, G., Perez-Riverol, Y., Reisinger, F., Ternent, T., et al. (2016). 2016 update of the PRIDE database and its related tools. Nucleic Acids Res 44, D447–456.

Wen, X., Tan, W., Westergard, T., Krishnamurthy, K., Markandaiah, S.S., Shi, Y., Lin, S., Shneider, N.A., Monaghan, J., Pandey, U.B., et al. (2014). Antisense proline-arginine RAN dipeptides linked to C9ORF72-ALS/FTD form toxic nuclear aggregates that initiate in vitro and in vivo neuronal death. Neuron 84, 1213–1225.

Westergard, T., Jensen, B.K., Wen, X., Cai, J., Kropf, E., Iacovitti, L., Pasinelli, P., and Trotti, D. (2016). Cell-to-Cell Transmission of Dipeptide Repeat Proteins Linked to C9orf72-ALS/FTD. Cell Rep 17, 645–652.

White, M.R., Mitrea, D.M., Zhang, P., Stanley, C.B., Cassidy, D.E., Nourse, A., Phillips, A.H., Tolbert, M., Taylor, J.P., and Kriwacki, R.W. (2019). C9orf72 Poly(PR) Dipeptide Repeats Disturb Biomolecular Phase Separation and Disrupt Nucleolar Function. Mol Cell 74, 713–728 e716.

Wood, M.J.A., Talbot, K., and Bowerman, M. (2017). Spinal muscular atrophy: antisense oligonucleotide therapy opens the door to an integrated therapeutic landscape. Hum Mol Genet 26, R151–R159.

Yin, S., Lopez-Gonzalez, R., Kunz, R.C., Gangopadhyay, J., Borufka, C., Gygi, S.P., Gao, F.B., and Reed, R. (2017). Evidence that C9ORF72 Dipeptide Repeat Proteins Associate with U2 snRNP to Cause Mis-splicing in ALS/FTD Patients. Cell Rep 19, 2244–2256.

Zhang, Y.J., Gendron, T.F., Ebbert, M.T.W., O’Raw, A.D., Yue, M., Jansen-West, K., Zhang, X., Prudencio, M., Chew, J., Cook, C.N., et al. (2018). Poly(GR) impairs protein translation and stress granule dynamics in C9orf72-associated frontotemporal dementia and amyotrophic lateral sclerosis. Nat Med.

Zhang, Y.J., Guo, L., Gonzales, P.K., Gendron, T.F., Wu, Y., Jansen-West, K., O’Raw, A.D., Pickles, S.R., Prudencio, M., Carlomagno, Y., et al. (2019). Heterochromatin anomalies and double-stranded RNA accumulation underlie C9orf72 poly(PR) toxicity. Science 363.

Zhou, Q., Lehmer, C., Michaelsen, M., Mori, K., Alterauge, D., Baumjohann, D., Schludi, M.H., Greiling, J., Farny, D., Flatley, A., et al. (2017). Antibodies inhibit transmission and aggregation of C9orf72 poly-GA dipeptide repeat proteins. EMBO Mol Med 9, 687–702.

Zu, T., Liu, Y., Banez-Coronel, M., Reid, T., Pletnikova, O., Lewis, J., Miller, T.M., Harms, M.B., Falchook, A.E., Subramony, S.H., et al. (2013). RAN proteins and RNA foci from antisense transcripts in C9ORF72 ALS and frontotemporal dementia. Proc Natl Acad Sci U S A 110, E4968–4977.

