## Supplemental Figures 1-4; Tables 1,2 for "Generalized displacement of DNA- and RNA-binding factors mediates the toxicity of arginine-rich cell-penetrating peptides"

Lafarga *et al* **Fig S1**
(related to **Fig 1**)

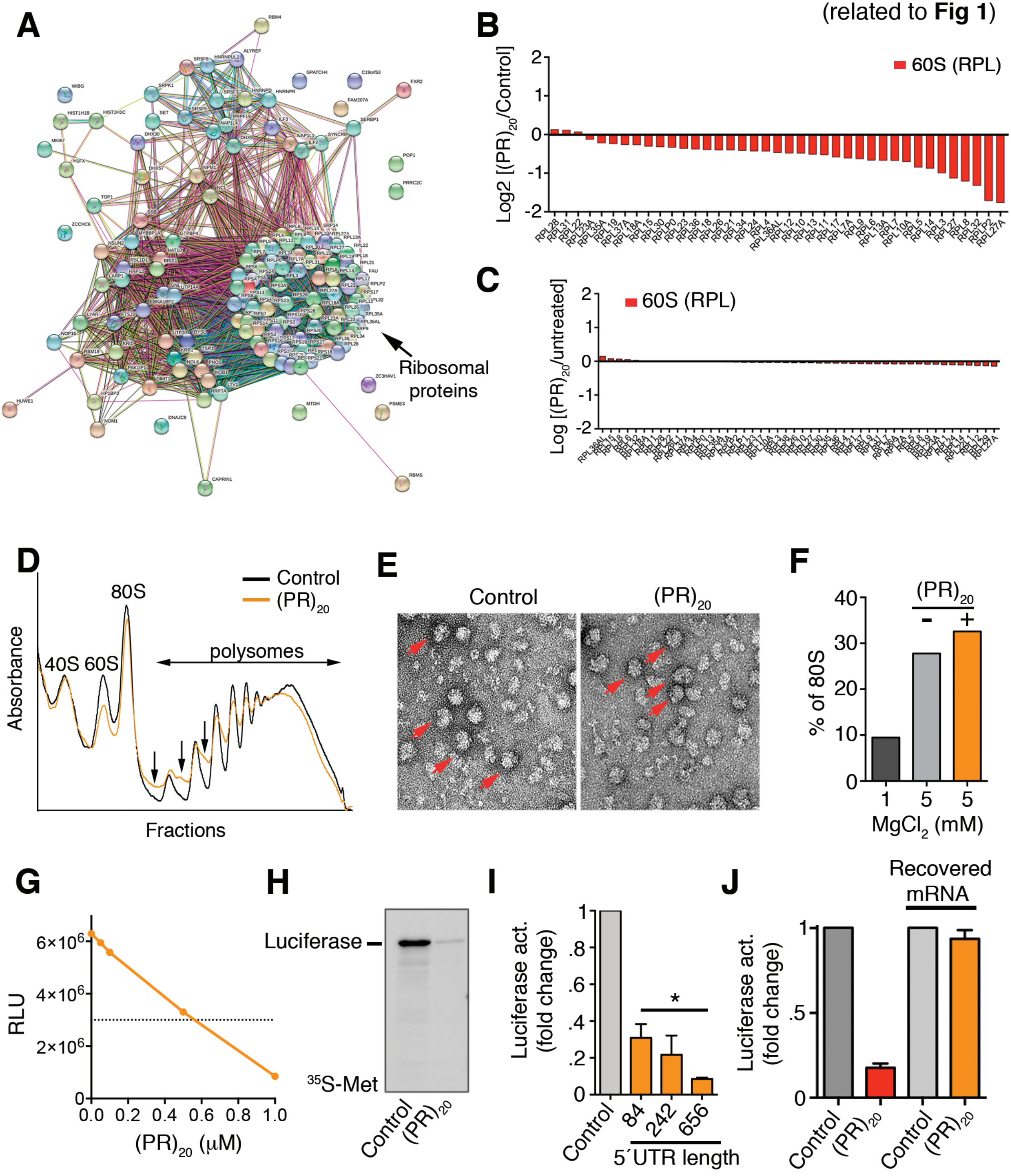

**Fig. S1.** (Related to **Fig. 1**) (**A**) STRING analysis (https://string-db.org) of the protein interaction networks found among the proteins identified in the purification of ribosomes from RPS9$^{SBP}$-expressing HeLa cells. The node containing 60S and 40S factors (RPLs and RPSs) is indicated (arrow). The panel illustrates the ribosome composition from $(PR)_{20}$-untreated cells. (**B**) Protein levels of RPL factors in ribosomes purified from Hela RPS9$^{SBP}$ cells exposed to 10 μM of $(PR)_{20}$ for 16h, as identified by LC-MS/MS. (**C**) Protein levels of RPL factors in the input extracts used for ribosome purification from Hela RPS9$^{SBP}$ cells exposed to 10 μM of $(PR)_{20}$ for 16h, as identified by LC-MS/MS. (**D**) Representative polysome profiles obtained from HeLa cells untreated or treated with 10 μM of (PR)20 for 16h. The presence of halfmers is indicated (arrows). (**E**) Electron microscopy images from purified 40S and 60S ribosomal complexes (1 pmol each) assembled *in vitro* in the presence of $MgCl_2$, and in the presence or absence of 5 pmol of $(PR)_{20}$. Assembled 80S particles are indicated (red arrows). (**F**) Quantification of 80S particles identified in (**E**) (n=1000) in non-assembly (1 mM $MgCl_2$) or assembly (5 mM $MgCl_2$) conditions. (**G**) *In vitro* translation of 100 ng of luciferase mRNA (quantified by Relative Luciferase Units (RLU)) in the presence of increasing doses of $(PR)_{20}$. (**H**) *In vitro* translation of 100 ng of luciferase mRNA in the presence or absence of 0.5 μM $(PR)_{20}$. Translation products were labeled with [$^{35}$S]-Met/Cys and analyzed by SDS-PAGE and autorradiography. (**I**) *In vitro* translation of 100 ng of luciferase mRNA with different 5' UTR lengths in the presence or absence of 0.5 μM $(PR)_{20}$. (**J**) *In vitro* translation of 100 ng of luciferase mRNA in the presence or absence of 0.5 μM $(PR)_{20}$. In the right two columns, the mRNA was extracted from a translation reaction done in the presence of the DPR, and subsequently used in a new translation reaction performed in the absence of $(PR)_{20}$. *, $p<0.05$.

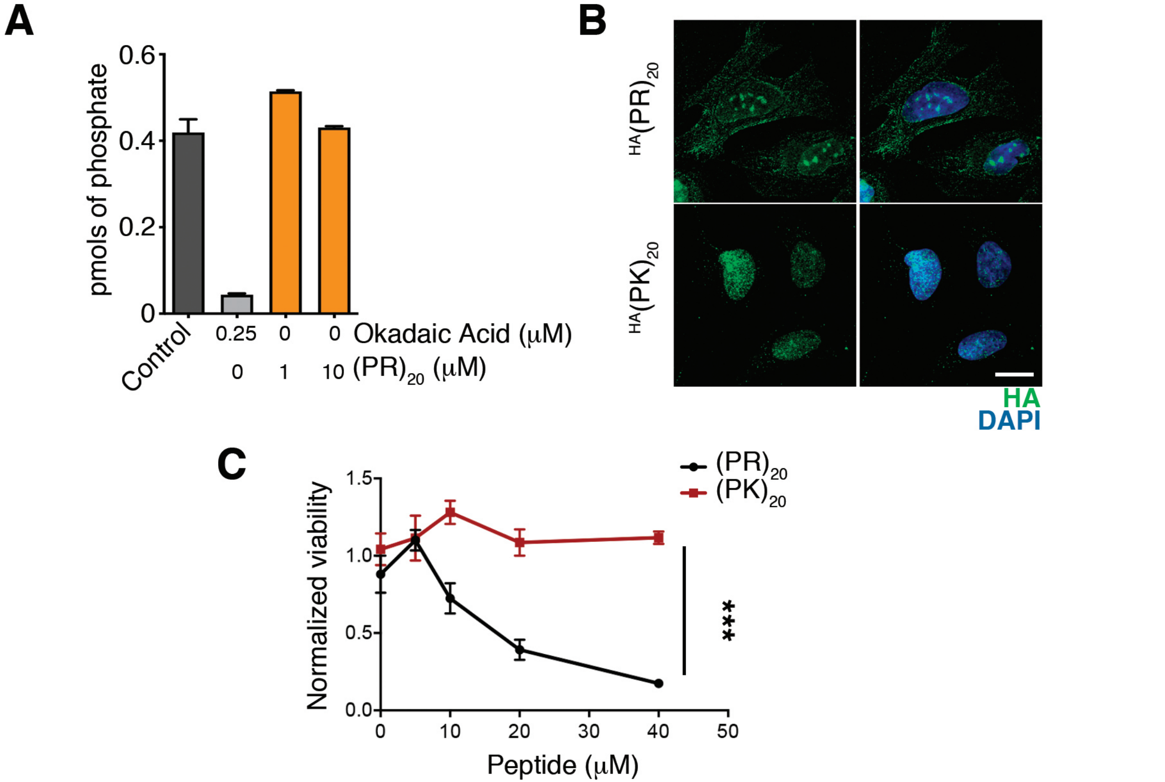
A
pmols of phosphate
0.6
0.4
0.2
0
Control
0.25
0
0
Okadaic Acid (μM)
0
1
10
(PR)20 (μM)
B
HA(PR)20
HA(PK)20
HA
DAPI
C
Normalized viability
1.5
1.0
0.5
0.0
0
10
20
30
40
50
Peptide (μM)
(PR)20
(PK)20
***

**Figure S2** (*related to* ***Fig. 4***). (**A**) A PP2A in vitro phosphatase assay performed in the presence or absence of $(PR)_{20}$ or Okadaic acid (a phosphatase inhibitor). PP2A activity is measured through the liberation of phosphates from a KRpTIRR peptide. (**B**) Immunofluorescence of HA-$(PR)_{20}$ or $(PK)_{20}$ (green) in U2OS cells treated with 7.5 μM of each peptide for 1 h. Scale bar (white) indicates 2.5 μm. DAPI was used to stain DNA. (**C**) Percentage of viable cells as evaluated with a CellTiter-Glo luminescent assay in U2OS cells treated with increasing doses of $(PR)_{20}$ or $(PK)_{20}$ peptides at the indicated doses. ****, $p<0.0001$

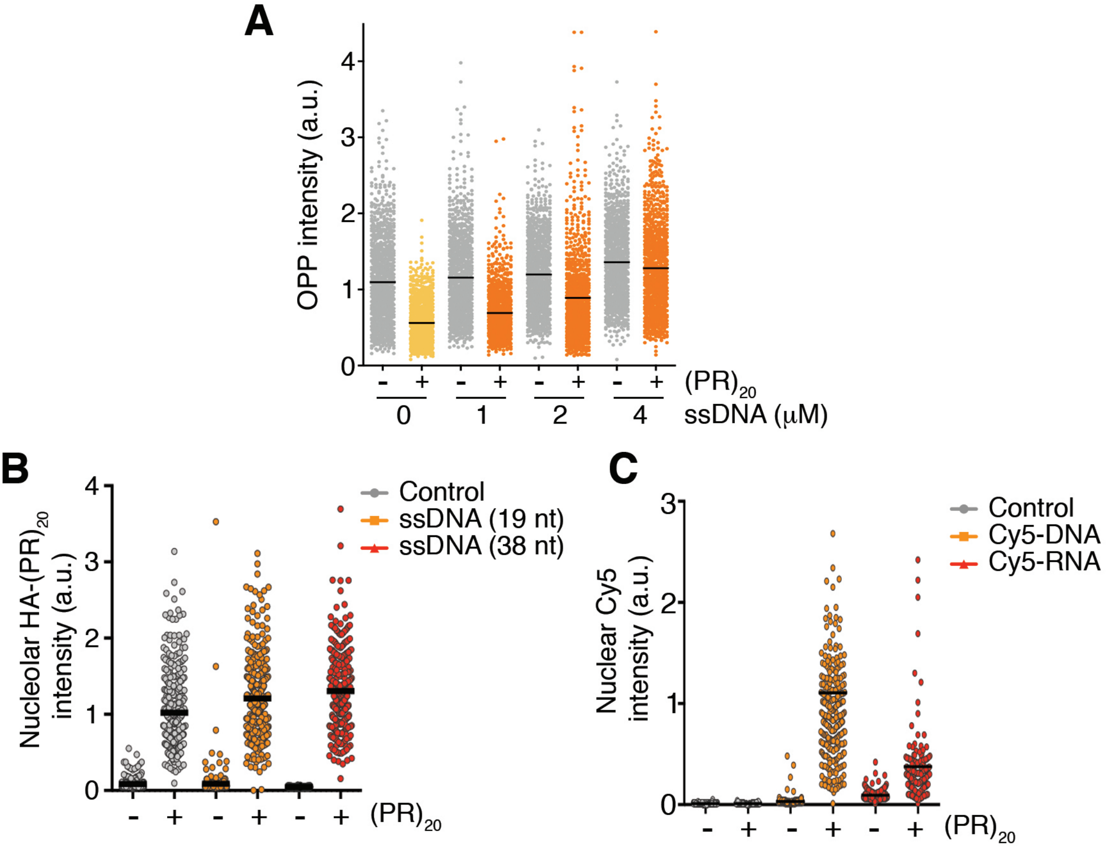
A
OPP intensity (a.u.)
0
1
2
3
4
- + - + - + - +
(PR)20
0 1 2 4
ssDNA (μM)
B
Nucleolar HA-(PR)20 intensity (a.u.)
0
1
2
3
4
Control
ssDNA (19 nt)
ssDNA (38 nt)
- + - + - +
(PR)20
C
Nuclear Cy5 intensity (a.u.)
0
1
2
3
Control
Cy5-DNA
Cy5-RNA
- + - + - +
(PR)20

**Figure S3** (*related to Fig. 3*). (**A**) HTM-mediated quantification of OPP levels per cell $(PR)_{20}$ (10 μM, 16h) alone or together with a 38 nt ssDNA oligonucleotide at the indicated doses. (**B**) HTM-mediated quantification of the nucleolar levels of a HA-$(PR)_{20}$ peptide in U2OS cells treated with 7.5 μM HA-$(PR)_{20}$ alone or together with 2 μM of 19 or 38 nt ssDNA oligonucleotides for 8 h. (**C**) HTM-mediated quantification of the nuclear levels of Cy5-labeled ssDNA or ssRNA 19 nt oligonucleotides in U2OS cells treated with 7.5 μM HA-$(PR)_{20}$ alone or together with 4 μM of the oligonucleotides for 8 h. Black lines indicate mean values. Examples of the images used for the analysis shown in (**B, C**) are shown in **Fig. 3D, E**.

Lafarga *et al* **Fig S4**
(related to **Fig 4**)

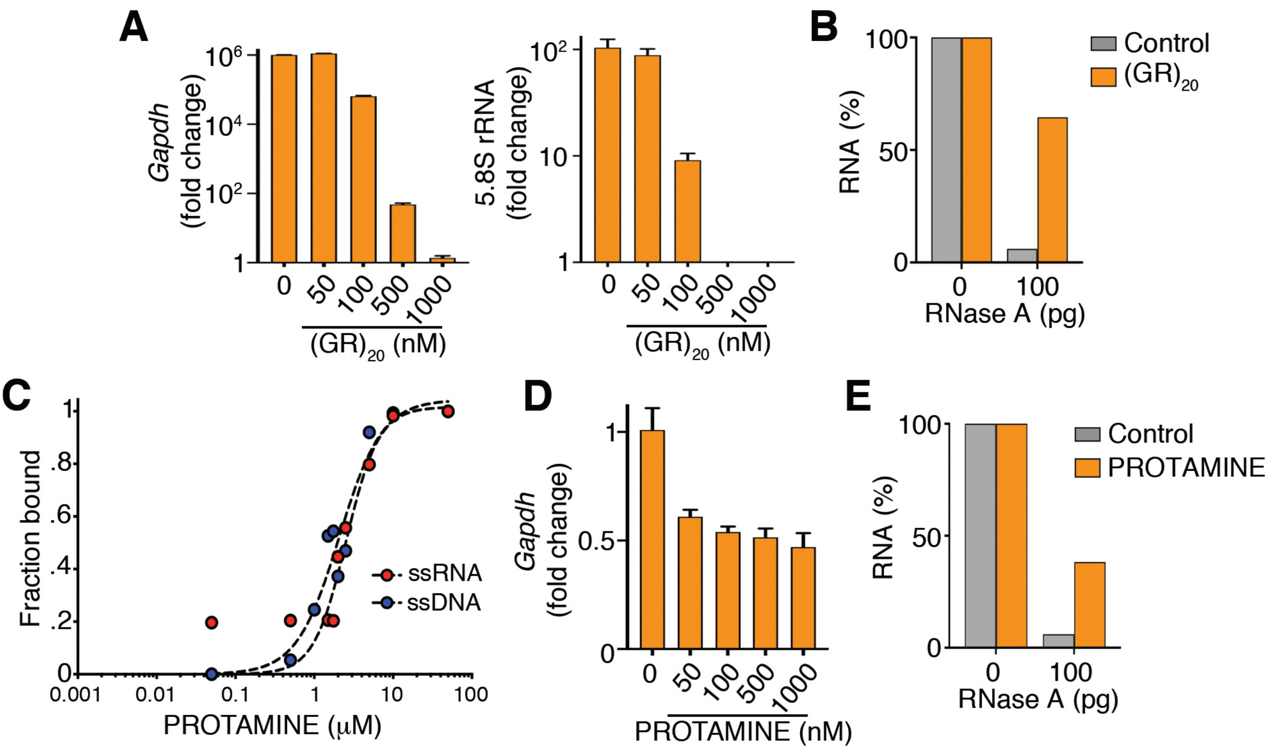


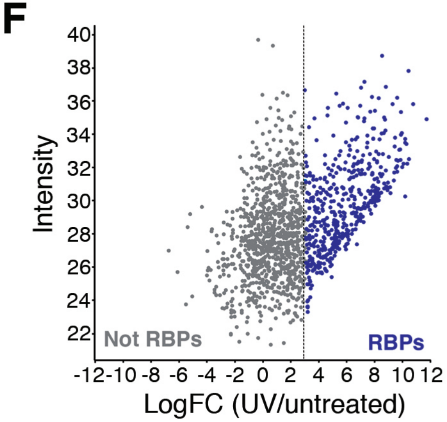


**G**

| | **Biological Process (GO)** | | |
|---|---|---|---|
| Pathway ID | *Pathway description* | *Count in gene set* | *False Discovery Rate* |
| *GO:0006396* | RNA processing | 204 | 5.96 e-154 |
| *GO:0016071* | mRNA metabolic process | 180 | 2.45 e-141 |
| *GO:0022613* | Ribonucleoprotein complex biogenesis | 126 | 6.10 e-112 |
| *GO:0016070* | RNA metabolic process | 315 | 3.18 e-98 |
| *GO:0010467* | Gene expression | 333 | 1.04 e-97 |

*Pathway analysis on the **RBP** fraction*

**Figure S4** (*related to* ***Fig. 4***). (**A**) Percentage of *GAPDH* or 5.8 rRNA levels quantified by qPCR in reactions containing increasing doses of $(GR)_{20}$ (n=3). (**B**) Percentage of RNA (1 μg) remaining after a 15´digestion with 100 pg of RNase A in the presence or absence of $(PR)_{20}$ (5 μM). (**C**) Quantification of EMSA assays evaluating the binding of PROTAMINE to 19 nt-long ssDNA and ssRNA molecules. Each probe (0.2 μM) was incubated with increasing concentrations of $(PR)_{20}$ for 10´. Curve-fitting was performed using non-linear regression with the Hill equation. (**D**) Percentage of *GAPDH* levels quantified by qPCR in reactions containing increasing doses of PROTAMINE (n=3). (**E**) Percentage of RNA (1 μg) remaining after a 15´digestion with 100 pg of RNase A in the presence or absence of PROTAMINE (5 μM). (**F**) Distribution of the proteins and their levels from the proteomic analysis following isolation of mRNA-binding proteins. Proteins that showed significant enrichment on the olido(dT)-captured fractions following UV crosslinking were identified as RBPs for subsequent analyses. (**G**) Gene Ontology (GO) analysis illustrating the biological pathways that were most significantly enriched among the proteins defined as RBPs in the proteomic experiment shown in (**F**). Note that, indeed, all biological pathways are related to RNA metabolism.

**Table S1.** Proteins with significantly reduced levels on ribosomal fractions purified from $(PR)_{20}$-treated HeLa-RPS9$^{SBP}$ cells. ***Related to Fig 1***.

| Gene Name | $Log_2$ ($(PR)_{20}$/control) |
|---|---:|
| PSME3 | -3.61 |
| H1FX | -1.94 |
| RPL27A | -1.76 |
| RPLP2 | -1.71 |
| BRIX1 | -1.52 |
| GTPBP4 | -1.35 |
| RPL32 | -1.32 |
| KRR1 | -1.22 |
| NAP1L4 | -1.21 |
| HP1BP3 | -1.20 |
| RPL3 | -0.99 |
| RPL14 | -0.87 |
| RPL10A | -0.70 |
| RPL6 | -0.66 |
| UTP14A | -0.65 |
| RPL9 | -0.63 |

**Table S2.** Proteins that show statistically significant reduced levels on chromatin after treatment of U2OS cells with $(PR)_{20}$ (20 μM) or PROTAMINE (30 μM). *Related to* ***Figure 4.***

| Gen Symbol | $Log_2((PR)_{20}/control)$ | $Log_2(PROTAMINE/control)$ |
|---|---:|---:|
| FURIN | -2.45 | -3.61 |
| POTEJ | -2.05 | -2.15 |
| POTEKP | -1.72 | -1.63 |
| TMEM126B | -1.40 | -1.19 |
| POTEF | -1.38 | -1.20 |
| HIST1H1C | -1.33 | -0.61 |
| FARS2 | -1.28 | -0.75 |
| HIST1H1A | -1.27 | -0.46 |
| SNX5 | -1.06 | -0.82 |
| H1F0 | -1.05 | -0.39 |
| PITPNA | -1.02 | -0.60 |
| PDK2 | -1.01 | -0.64 |
| GLTSCR2 | -1.01 | -0.42 |
| EEF1B2 | -1.01 | -0.86 |
| EEF1G | -0.98 | -0.75 |
| GAPDH | -0.97 | -0.80 |
| NRF1 | -0.96 | -0.90 |
| PSMF1 | -0.93 | -0.42 |
| AURKC | -0.92 | -0.87 |
| CORO1B | -0.88 | -0.47 |
| EBAG9 | -0.87 | -0.64 |
| APEX1 | -0.86 | -0.70 |
| EEF1D | -0.83 | -0.69 |
| ALDOC | -0.82 | -0.45 |
| REPIN1 | -0.82 | -0.54 |
| H2AFY | -0.82 | -0.43 |
| RXRB | -0.75 | -0.50 |
| HMGA1 | -0.75 | -0.68 |
| TPT1 | -0.73 | -0.65 |
| RPUSD4 | -0.72 | -0.87 |
| DNAJC15 | -0.70 | -0.40 |
| RING1 | -0.70 | -0.57 |
| PCBP3 | -0.67 | -0.41 |
| HMGA2 | -0.66 | -0.51 |
| SLC25A40 | -0.65 | -0.41 |
| HN1L | -0.65 | -0.40 |
| MEN1 | -0.65 | -0.40 |
| MSN | -0.64 | -0.58 |
| HDGF | -0.62 | -0.44 |
| HCFC1 | -0.62 | -0.52 |
| TGIF2LX | -0.62 | -0.52 |
| HMGB1;HMGB1P1 | -0.62 | -0.50 |
| LRRC57 | -0.60 | -0.49 |
| FLNA | -0.60 | -0.50 |
| RANBP1 | -0.60 | -0.59 |
| NCKIPSD | -0.59 | -0.61 |

| | | |
|---|---|---|
| EZR | -0.58 | -0.51 |
| CMSS1 | -0.58 | -0.35 |
| LANCL2 | -0.58 | -0.58 |
| CSRP2 | -0.58 | -0.48 |
| TMF1 | -0.58 | -0.44 |
| EEF1A2 | -0.56 | -0.54 |
| TARS | -0.56 | -0.45 |
| CDYL | -0.56 | -0.75 |
| DIDO1 | -0.55 | -0.44 |
| AHNAK | -0.55 | -0.48 |
| ACAP2 | -0.55 | -0.39 |
| SYNGR3 | -0.55 | -0.60 |
| PALM2 | -0.55 | -0.48 |
| ARHGAP17 | -0.55 | -0.57 |
| ID1 | -0.55 | -0.39 |
| CCAR1 | -0.54 | -0.40 |
| RPL18 | -0.54 | -0.38 |
| COPS7A | -0.54 | -0.51 |
| CCDC50 | -0.53 | -0.53 |
| ZBTB10 | -0.53 | -0.88 |
| CCNC | -0.53 | -0.38 |
| SFSWAP | -0.52 | -0.60 |
| SLC43A3 | -0.52 | -0.45 |
| WASF2 | -0.52 | -0.61 |
| SPR | -0.52 | -0.74 |
| GAP43 | -0.51 | -0.97 |
| FSCN1 | -0.51 | -0.40 |
| VPS39 | -0.51 | -0.57 |
| FUS | -0.50 | -0.46 |
| SARS | -0.50 | -0.46 |
| FIP1L1 | -0.49 | -0.38 |
| DIAPH2 | -0.49 | -0.67 |
| CSTF2 | -0.49 | -0.40 |
| SCAF4 | -0.49 | -0.36 |
| CBX8 | -0.48 | -0.47 |
| PRPF38B | -0.48 | -0.35 |
| DCUN1D1 | -0.47 | -0.61 |
| DDX31 | -0.47 | -0.45 |
| PDLIM7 | -0.47 | -0.41 |
| RECQL | -0.47 | -0.36 |
| FNBP4 | -0.47 | -0.40 |
| EHD3 | -0.46 | -0.38 |
| SCAF1 | -0.46 | -0.37 |
| MRGBP | -0.46 | -0.54 |
| MAVS | -0.45 | -0.38 |
| CORO1C | -0.45 | -0.55 |
| MINA | -0.45 | -0.48 |
| COPS8 | -0.45 | -0.74 |
| TCERG1 | -0.45 | -0.42 |
| NEDD1 | -0.44 | -0.59 |
| SEPT9 | -0.44 | -0.43 |
| HDGFRP3 | -0.44 | -0.36 |
| FAHD1 | -0.44 | -0.73 |

| | | |
|---|---|---|
| FKBP3 | -0.43 | -0.42 |
| PA2G4 | -0.43 | -0.42 |
| DDX42 | -0.43 | -0.40 |
| CASK | -0.42 | -0.69 |
| GABPA | -0.41 | -0.36 |
| MAD2L1BP | -0.40 | -0.40 |
| ITPRIP | -0.40 | -0.67 |
| EHD4 | -0.40 | -0.36 |
| NARS | -0.39 | -0.41 |
| EEF1A1;EEF1A1P5 | -0.39 | -0.47 |
| WDR33 | -0.39 | -0.42 |
| DEK | -0.39 | -0.44 |
| TBP | -0.39 | -0.36 |
| DOK1 | -0.39 | -0.35 |
| IFIT2 | -0.39 | -0.72 |
| UBAP2L | -0.38 | -0.49 |
| SEPT11 | -0.38 | -0.36 |
| RSU1 | -0.37 | -0.38 |
| NACA | -0.37 | -0.51 |
| ORC1 | -0.37 | -0.42 |
| CCNL1 | -0.37 | -0.37 |
| CEP97 | -0.37 | -0.45 |
| UBE2I | -0.36 | -0.42 |
| VPS13C | -0.36 | -0.41 |
| PRDX1 | -0.36 | -0.40 |
| DR1 | -0.36 | -0.64 |
| BCL7B | -0.35 | -0.36 |
| KDM2A | -0.35 | -0.41 |
